# How to use transcriptomic data for game-theoretic modeling of treatment-induced resistance in cancer cells? A case study in patient-derived glioblastoma organoids

**DOI:** 10.1101/2022.01.26.477755

**Authors:** Louise Spekking, Christer Lohk, Weronika Jung, Maikel Verduin, Sepinoud Azimi, Christopher Hubert, Marc Vooijs, Rachel Cavill, Kateřina Staňková

**Affiliations:** Delft University of Technology; Bordeaux University; Maastricht University; Cleveland Clinic

**Keywords:** Qualitative resistance, evolutionary dynamics, game theory, transcriptomics, glioblastoma, patient-derived organoids, radioresistance

## Abstract

Game theory is a powerful tool to model strategic decision making, but also interactions within Darwinian biological systems, such as cancer. As such, in the past decades, game-theoretical models have helped to understand cancer, its response to various treatments, and to design better therapies. However, to fully utilize the potential of game-theoretical modelling in designing better anti-cancer therapies, we need more information on cancer population (ecological) and strategy (evolutionary) dynamics in response to treatment for each patient and their tumors. Here we explore how transcriptomics data can be utilized as an input of game-theoretical models for predicting evolutionary response to irradiation in patient-derived glioblastoma organoids. For that purpose, we utilize both supervised and unsupervised machine learning methods to identify relevant cancer cell types and how their proportions change over time in the organoids. We then fit these proportions to the replicator dynamics, the most common evolutionary game dynamics, to predict both transient evolutionary dynamics and evolutionary stable strategy (ESS) cell proportions. Our predictions in glioblastoma organoids suggest that hypoxia is the most important factor in identifying short-term response to irradiation, while this seems much less relevant for the long-term response corresponding to the ESSs. Further, we conclude that supervised methods are the best way to estimate cancer evolutionary dynamics when therapy resistance is a qualitative evolutionary trait. We believe that our methodology can help in designing better therapies through testing evolutionary responses in patient-derived organoids, while in parallel the ecological response can be tracked through serum biomarkers and imaging in the corresponding patients.

## 1 Introduction & Motivation

One of the biggest obstacles in treating advanced cancers is therapy resistance [1, 2, 3, 4, 5, 6, 7, 8]. Game-theoretic models of cancer and its treatment led to first evolutionary therapies (also known as adaptive or Darwinian therapies), i.e. therapies anticipating and preventing the evolution of therapy resistance in cancer cells [9, 10, 11, 12, 13, 14, 15, 16, 17, 18]. Ongoing clinical trials on evolutionary therapies (such as NCT02415621, NCT03511196, NCT03630120, NCT03543969, NCT05393791 (ANZadapt), NCT05080556, NCT04343365) show very promising results in terms of quality and quantity of patients’ life.

In Zhang et al’s metastatic castrate-resistant prostate cancer trial, the prostate-specific antigen (PSA) was used as a proxy for the tumor size in the course of the treatment [19]. Because of the lack of information on the composition of the tumors, possible initial tumor compositions were estimated from in-vitro measurements and expected interactions among cancer cells of different types [20, 9, 21]. This has led to 22 different possible parametrizations of the model, differing in their predictions regarding proportion of therapy resistant cells in the evolutionary stable equilibrium. These 22 cases were clustered into 3 different groups: best-responder, responder, and non-responder [20, 9, 22, 23]. In simulations, Zhang et al’s protocol was shown to be more effective than the maximum tolerable dose corresponding to the standard of care for all these parametrizations [9, 22, 23, 24]. This was confirmed in the trial itself [19, 21]. In the absence of detailed information on tumor composition, the treatment protocol was designed such that it was effective for all possible initial tumor compositions. These different compositions had to be considered as tumors with the same size but different composition will likely respond to the same treatment differently [6, 24]. More precise information on the tumor composition in the course of treatment is needed to design even more effective evolutionary therapies [25, 2]. A rough estimate of tumor composition may be made from cancer cells’ response to initial treatment application, as proposed in [26]. Another way, explored in the current study, is to utilize transcriptomics data from patient-derived organoids to model and predict how the proportions of different cancer cell types change over time in response to the treatment [27, 28, 29].

Ideally, single cell technologies would give us this information, however these techniques are prohibitively expensive, both in monetary terms and in terms of the computational and expert bioinformaticians’ time that are needed to deal with the vast amounts of data these techniques produce [30]. Despite this, upfront costs are dropping and some studies have been conducted using this technology to study particular cancers e.g. [31]. If costs either in processing time/effort or money were to remain high this would prevent their routine use in the clinic, especially in scenarios where patients need long term regular monitoring to track the status of their tumor.

Therefore, here we will focus on techniques to obtain this information from bulk sequencing data, where data is obtained from many cells simultaneously using standard omics techniques, such as transcriptomics [32]. These deconvolution methods come in two distinct classes, supervised - where the training set is furnished with the known proportions of the cell types, or the profile of each cell type is known in advance and unsupervised - where nothing is assumed about the proportions nor the profiles of the cell types in advance of the analysis [33]. In this paper we test a range of approaches from each class.

We compare supervised and unsupervised methods on the data from patient-derived glioblastoma organoids to obtain cell profiles and proportions for the cell types present in glioblastoma [29, 27]. The organoids provide a platform on which we can test potential treatments and observe the treatment effects *in vitro*. With organoids, where the total tumor cell population is kept constant, we can track how treatment impacts the cell type composition over time without needing invasive biopsies. The tumor burden change over time can be tracked in the patient blood biomarker and imaging data. This makes organoids an ideal testing ground for the game-theoretic models. After identifying different cell types in organoids and how their proportions change over time in response to radiotherapy, we fit the most classical game-theoretic model, replicator dynamics, to this time series data. We hereby provide a methodology to utilize bulk data for estimating cancer evolutionary dynamics, which will then lead to better evolutionary therapies.

## 2 Methods

### 2.1 Datasets

In this work we make use of several datasets, which we describe below.

#### 2.1.1 TGCA Glioblastoma dataset

The Cancer Genome Atlas (TCGA) is a publicly available resource containing molecular data for over 10000 specimens, from at least 33 different types of cancer and metadata linking this information to clinical data for the patients [34]. We used the TCGA-GBM project, which focuses on glioblastoma samples. We used 168 samples with available RNA-SEQ data. For each sample we retrieved the FPKM-UQ normalised RNA-SEQ. These files provide measurements for over 60000 RNA sequences per sample.

#### 2.1.2 Neftel’s single-cell data

We utilized Neftel et al.’s single-cell dataset acquired through full-length scRNA-SEQ (Smart-seq2) [35]. We downloaded this dataset from NIH Gene Expression Omnibus (GEO: GSE131928). It contains preprocessed single cell RNA-seq expression profiles of adult and paediatric IDH-wildtype Glioblastomas from 28 tumors with an average of 5730 genes detected per cell. From the initial 5730 genes, 289 were selected for cell type distribution analysis based on meta-module signatures. scRNA-SEQ data were complemented with metadata from Broad Institute Single Cell Portal (Study: Single cell RNA-seq of adult and pediatric glioblastoma) containing meta-module (NPC1, NPC2, MES1, MES2, OPC, AC) scores for 7930 cells. Cells were categorized according to their highest corresponding meta-module scores.

#### 2.1.3 Organoid data

The patient-derived organoids were grown as per the protocol in [29] and then treated with either 4 Gy or 10 Gy irradiation. Samples were collected pre-treatment, directly post-treatment, and at 24 and 72 hours post-treatment. RNA was isolated using Nucleospin RNA (Macherey-Nagel) according to the manufacturers protocol. RNA concentration and integrity was confirmed using 2100 Bioanalyzer (Agilent Technologies) . RNA was hybridized to affymetrix human microarrays by Eurofins (Denmark). All conditions were tested in GBM organoids derived from 3 patients.

### 2.2 Unsupervised methods

For the unsupervised methods, we focus on non-negative matrix factorisation (NMF), a method which describes the data matrix as a product of two matrices, one describing the cell type proportions and the other describing the profiles of each cell type. This assumes that the data can be approximated simply by taking a weighted sum of the cell type profiles where the weights are the proportions of each cell type present [36].

The deconvolution of cancer cells can be seen as a specific case of the blind source separation (BSS) issue [37], i.e. separating a group of source signals from measurements of mixed signals. A cancer tumor, as a heterogeneous mixture of different cell types, can be analysed using a linear BSS model, as this model does not require knowledge of any predefined cell type signatures and can therefore be used to formulate the deconvolution problem. Whilst there are various methods which have been applied for unsupervised deconvolution of bulk cell profiles [38, 39, 40], in earlier unpublished work we showed that Non-Negative matrix factorisation (NMF) is by far the most promising approach that agrees with the literature findings [37, 40]. Therefore, here we focused on the family of non-negative matrix factorisation-based approaches. In this study we applied several NMF algorithms [41, 42, 43, 38, 44]. The number of cell types selected should correspond to the NMF rank for which the best results are obtained. To evaluate the obtained results, we used several classic indicators (Cophenetic coefficient [41], Dispersion coefficient [44], Residual Sum of Squares [45], Silhouette Coefficient [46]), whose values indicated the optimal factorisation rank.

### 2.3 Supervised methods

For the supervised approaches we use Scaden (Single-cell Assisted Deconvolutional Network), a reference-based deconvolution framework to determine the cellular composition of glioblastoma patient-derived organoids from microarray profiles [47]. As a reference, we rely on single-cell profiles and a blueprint suggested by Neftel et al. that assigns cells into four meta-modules, each characterized by a unique gene expression signature and genetic alterations [35].

Scaden is a deep learning framework that utilizes scRNA-SEQ expression data to train a model for the approximation of cell-type proportions from an input bulk expression profile [47]. The training requires vast amounts of bulk RNA-SEQ expression profiles with corresponding cell type proportions as ground truth. This was achieved through random subsampling of 6750 single-cell profiles from Neftel’s scRNA-SEQ dataset into 1000 pseudo-bulk RNA-SEQ gene expression profiles containing sum of expression values by gene. After the training set generation, genes with zero expression across all samples or with variance less than 0.1 were removed from bulk expression and training profiles. Furthermore, only genes that are present in both, training and inference datasets were used by the model. Next, the input data (both bulk expression profiles and training data) were transformed to the logarithmic space and scaled between 0 and 1. The model consists of three independent neural networks which are separately trained with preprocessed training data for 5000 training steps. The training was followed by inference to approximate cell type proportions from input bulk expression profiles (in addition to bulk RNA-SEQ, Menden et al. [47]) demonstrated reliable performance with a microarray dataset. The predictions for cell-type proportions were acquired from all three networks and averaged for the final output.

### 2.4 Game-theoretic model of glioblastoma

Changes in population characteristics such as size and density of cancer cells are associated with cancer ecological dynamics, while changes in heritable cell lineage characteristics (quantitative traits) or frequency of cancer cell types (qualitative traits) are considered evolutionary dynamics. This eco-evolution in the tumor can be captured with a use of various mathematical models. As the total cell population in organoids remains constant through the experiment, we analyze only changes in evolutionary dynamics. We utilize one of the most general approaches to integrate time scales of evolutionary dynamics; replicator dynamics [48, 49], which is the most classic game-theoretic dynamics used in modelling many biological processes as well as selection in population genetics. Given a finite set of cancer strategies *𝒯*, here understood as a set of cell types, their fitness matrix and their proportions at time *t*, the dynamics of cells of the same type can be described by deriving the following dynamics:

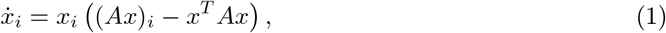

where *x*_*i*_ denotes the proportion of type *i ∈ 𝒯*, and consequently the column vector of proportions of all types is denoted by **x**. The *n × n* fitness matrix *A* = (*a*_*ij*_) defines the fitness of the considered cell types. If the entries of this matrix are between 0 and 1, each value *a*_*ij*_ defines the probability that a focal cell of type *i ∈ 𝒯* will produce an offspring of its own type when interacting with a cell of type *j ∈ 𝒯* . We can calculate the so-called evolutionary stable strategies (ESSs) of *A*, which maximize the fitness of the entire population and are uninvadable by a rare mutant with a different strategy. These ESSs may correspond to the attractors of the replicator dynamics (1).

## 3 Results

### 3.1 Results of unsupervised methods

As explained in Section 2.2, when applying the BSS model to define a deconvolution problem there is no prerequisite on prior knowledge regarding cell type signatures. In the absence of this information, one has to start the deconvolution by determining the number of cell types in the cancer mixture and only then determine their profiles and proportions. Here we created several simulated data sets and tested them with different NMF algorithms [41, 42, 43, 38, 44]. We used measures detailed in Section 2.2 to evaluate performance of these algorithms. These indicators on the simulated dataset mainly pointed to the actual value of the factoring rank. To disentangle the tumor mixture in glioblastoma, determine the number of component cancer cell types and determine their profiles, we compared results of the different NMF algorithms on the TCGA collection 2.1.1. We concluded that the measured gene expression values consist of three clusters, and therefore we used rank 3 of the factorisation. Subsequently, we approximated gene profiles of the clusters found in the tumor mixture. We then identified the proportion of each subtype (designated 1, 2 and 3) in the mixture before treatment was applied and tracked their changes in time after irradiation.

The changes in the proportions observed while the total population of cancer cells remains constant led us to a hypothesis that there are three types of cells in the mixture: The first type is sensitive to treatment (proportion in the mixture decreases), the second type is treatment-resistant (proportion in the mixture increases) and the for the last one the proportions remains unchanged with treatment.

To test this for each cell type obtained through NMF, we calculated Log2FoldChange (LFC) with respect to the other two cell types (one expression value subtracted from the other) for every gene. We then defined a threshold (LFC *>* 2). If the LFC was above 2 for both cases (LFCs calculated with respect to the other cell types), the gene was assigned to the signature of the corresponding cell type. We found that type 1, type 2 and type 3 signatures consist of 408, 15566, and 987 genes, respectively.

Taking these signature lists we adopted an over-representation approach [50], looking for which KEGG pathways are over-represented in the list of genes each signature. The most significant pathways (adjusted *p <* 0.001) from the KEGG database are shown in Table 1. Here we can see that the significant pathways in type 1 cells are related to infections, indicating genes involved in immune processes are important for those cells, thus type 1 may be immune cells. The corresponding processes could be then radiation-induced immunogenic cell death [51]. For type 2 cells we see some cancer pathways and cell cycle, indicating that the genes defining the type 2 cell profile are related to the tumor, typically proliferative cancer cells. Finally, for type 3 cells we see that the neuroactive ligand-receptor interaction pathway is the most significant hit, indicating that we are finding genes relating to more normal neuronal cells.

**Table 1:**
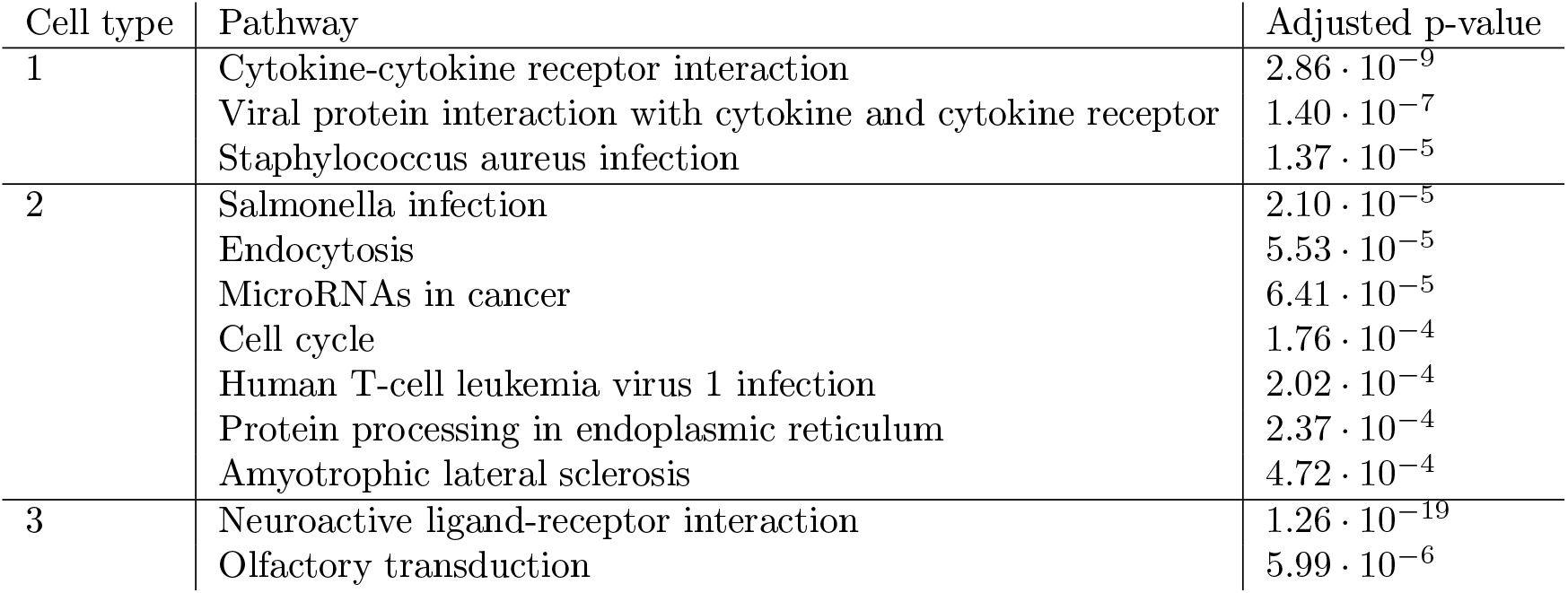
KEGG pathways with an adjusted *p <* 0.001 after over-representation analysis using ConsensusPathDataBase [50] with the genes from the NMF derived cell-type profiles and a background list of all genes used to generate the profiles.

As a further analysis we compared these genes to the genes from the meta-modules suggested by Neftel et al. We found that these were mostly covered by the type 2 cell type signature from the unsupervised model, as shown in Table 2, confirming as seen above that only one of the three types identified in our unsupervised analysis corresponds to proliferative tumor cells.

**Table 2:** Overlap of signature genes between Neftel’s meta-modules and cell types obtained through NMF (overlapping genes / total number of genes in the given Neftel’s meta-module). Signature genes for meta-modules have been made available by Neftel et al [35].

|  | MES1 | MES2 | AC | OPC | NPC1 | NPC2 | G1/S | G2/M |
| --- | --- | --- | --- | --- | --- | --- | --- | --- |
| V1 | 0/50 | 0/50 | 0/39 | 0/50 | 0/50 | 0/50 | 0/29 | 0/45 |
| V2 | 43/50 | 46/50 | 36/39 | 42/50 | 45/50 | 41/50 | 22/29 | 41/45 |
| V3 | 0/50 | 0/50 | 0/39 | 0/50 | 0/50 | 1/50 | 0/29 | 0/45 |

### 3.2 Results of supervised methods

In the supervised methods cell type proportions were inferred from microarray data using Scaden (see Section 2.3) The corresponding cell type proportions are shown for patient-derived organoids from three patients under two different conditions (4 and 10 Gy irradiation) both untreated and at 3 time points after treatment (Figure 1). For each patient, the first pre-exposure timepoint is shared between two conditions and the next three timepoints diverge due to differences in radiation doses. DNN-based model predictions indicate that more than 96% of all organoids (at all timepoints) consist of mesenchymal-like cells while the remaining 4% (or less) belong to other cell types present in Neftel’s dataset.

**Figure 1:**
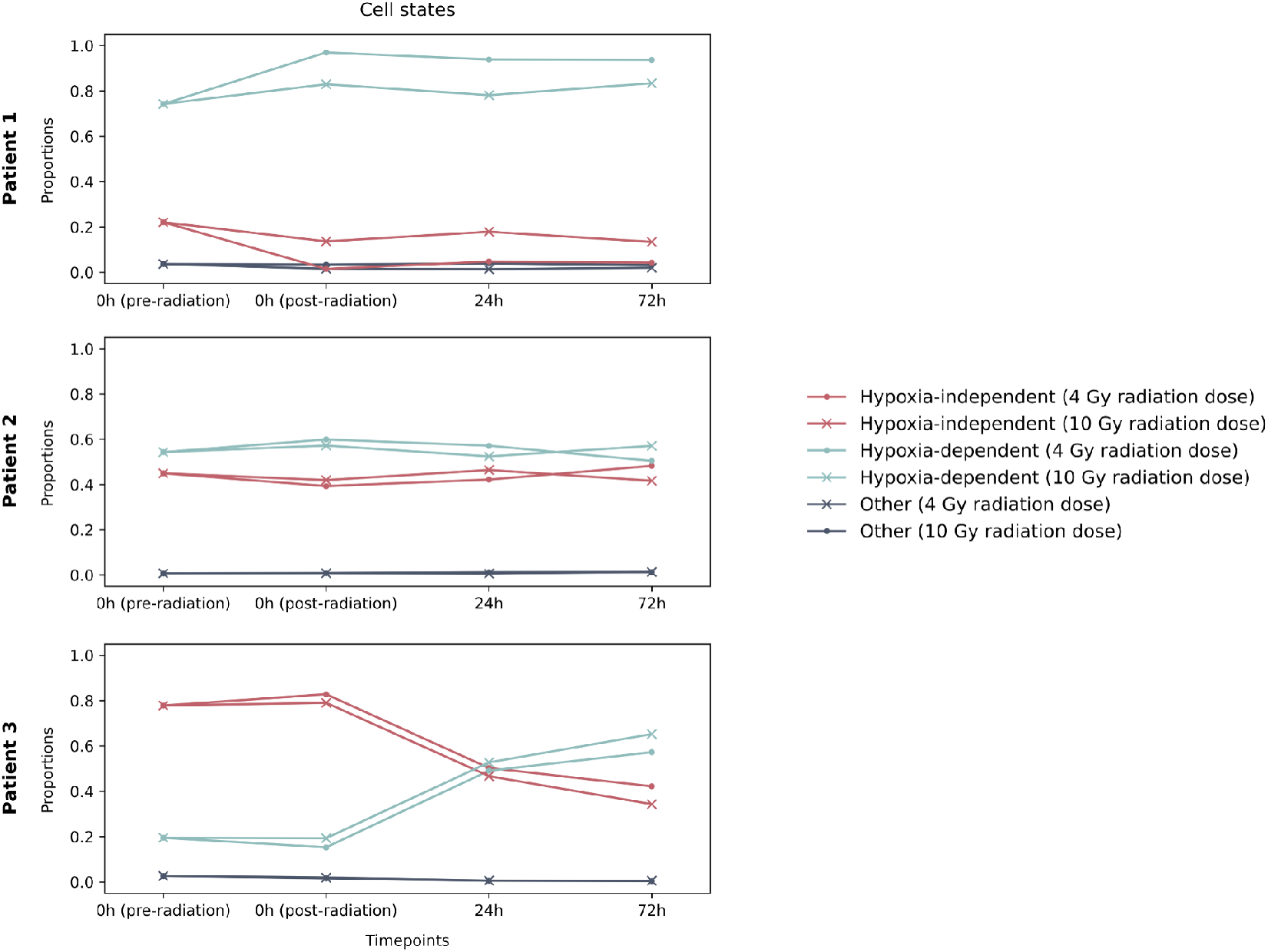
Population dynamics of mesenchymal-like and cycling cells in patient-derived glioma organoids under two divergent conditions. Hypoxia-independent cells are MES-like 1 and Hypoxia-depedent cells are MES-like 2 in Neftel’s scheme.

For patient 1 organoid, we observe a slight decline in the fraction of hypoxia-independent cells (MES-like 1) and an increase for hypoxia-dependent fraction post-radiation. Furthermore, a lower dose of radiation shows a greater effect on changes in cell-type proportions directly after the irradiation.

Patient 1’s organoid has the least varying proportions of different cell types.

The most visible changes in cell populations were recorded for patient 3. There is a clear trend of hypoxia-dependent cells (MES-like 2) overtaking hypoxia-independent cells (MES-like 1) as the primary cell type 72 hours post irradiation.

The dynamics in MES-like cell type could be caused by the upregulation of hypoxia-response genes in the hypoxia-independent cell population. This would lead to a transition from one cell type to another without any changes in population size. Similar dynamics could also be explained through different resistance to irradiation. If hypoxia-dependent cell types showed higher probability of survival (fitness) upon irradiation, as suggested by Wang et al. [52], we would observe a similar effect with a decrease in a total number of tumor cells due to apoptosis or other forms of cell death [53]. An alternative hypothesis revolves around the mechanism by which radiation works to cause DNA damage and cell death. This is achieved through the generations of reactive oxygen species, in hypoxic environments the necessary reactive oxygen can not be generated, so any cells surviving in hypoxic environments are naturally radiation resistant. This fits with previous observations that cells in the hypoxic core of glioblastoma organoids are resistant to radiation even when the cells in the normoxic environment at the edge of the organoid are not [29].

### 3.3 Results of game-theoretic model of organoids

As discussed earlier, comparison of the type profiles found by unsupervised methods with the meta-models proposed by Neftel et al. showed that most genes are covered by cell signatures from one of the three types. Thus, the assumption that the unsupervised methods result in different cancer cell types appears to be incorrect, hence the game theoretic model was only fitted to the results obtained by the supervised methods. The results obtained with the DNN model showed that organoids are mainly composed of mesenchymal cells. In order to simplify the tumor dynamics model, a replicator population was considered, which consists of two types of mesanchymal cells: hypoxia-dependent (MES-like 2) and hypoxia-independent (MES-like 1). Given two cell types in a population, the payoff matrix *A* has dimension 2*×*2, and its elements *a*_*ij*_ represent the probability of reproduction of cell type *i* when interacting with type *j*. As adding any constant to any column of the fitness matrix *A* does not change the dynamics, *a*_11_ and *a*_22_ are set to zero. To determine the true frequency dynamics defined by the fitness matrix, the remaining matrix parameters are fitted using available measurements in both of the environments created by treatment initialisation. The fitness matrix is assumed to have the following form:

**Table 3:** Fitness matrix *A* for cancer populations with two types of cells.

|  | MES 1 | MES 2 |
| --- | --- | --- |
| MES 1 | 0 | $a_{12}$ |
| MES 2 | $a_{21}$ | 0 |

For the found matrices, the residual sum of squares (RSS) between the available measurements and the fitted replicator dynamics is no more than 0.001 for each patient in both environments. The plots below 2 show the predicted dynamics of the two types (MES-like 1 and MES-like 2) and the relevant measurements. Horizontal lines in the figures represent evolutionary stable strategies (ESSs).

**Table 4:**
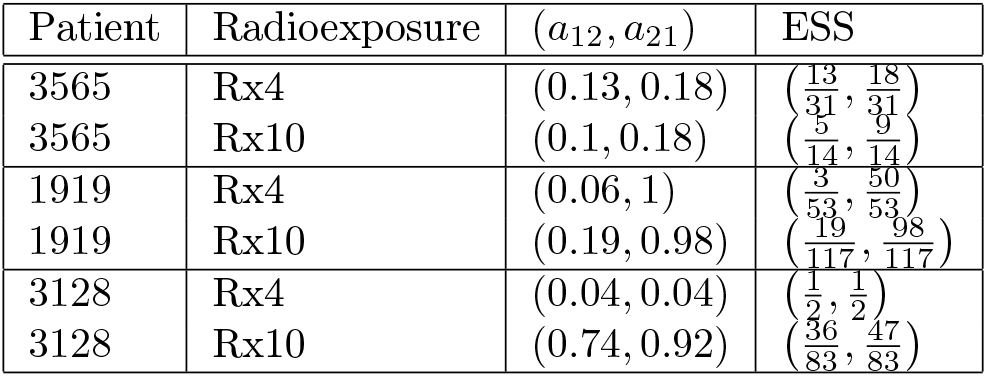
Fitted parameters of matrix *A* and calculated ESSs for the cancer population with two types.

| Patient | Radioexposure | $(a_{12}, a_{21})$ | ESS |
| --- | --- | --- | --- |
| 3565 | Rx4 | (0.13, 0.18) | $\left(\frac{13}{31}, \frac{18}{31}\right)$ |
| 3565 | Rx10 | (0.1, 0.18) | $\left(\frac{5}{14}, \frac{9}{14}\right)$ |
| 1919 | Rx4 | (0.06, 1) | $\left(\frac{3}{53}, \frac{50}{53}\right)$ |
| 1919 | Rx10 | (0.19, 0.98) | $\left(\frac{19}{117}, \frac{98}{117}\right)$ |
| 3128 | Rx4 | (0.04, 0.04) | $\left(\frac{1}{2}, \frac{1}{2}\right)$ |
| 3128 | Rx10 | (0.74, 0.92) | $\left(\frac{36}{83}, \frac{47}{83}\right)$ |

One of the characteristic features of advanced tumors is hypoxia, which is closely related to tumor radioresistance, as it can promote tumor progression and lead to poor clinical outcomes. In each patient, regardless of radiotherapy dose, the fitted model predicts in the long-term a majority of hypoxia-dependent cells, while both observed cancer cell types coexist at equilibrium.

**Figure 2:**
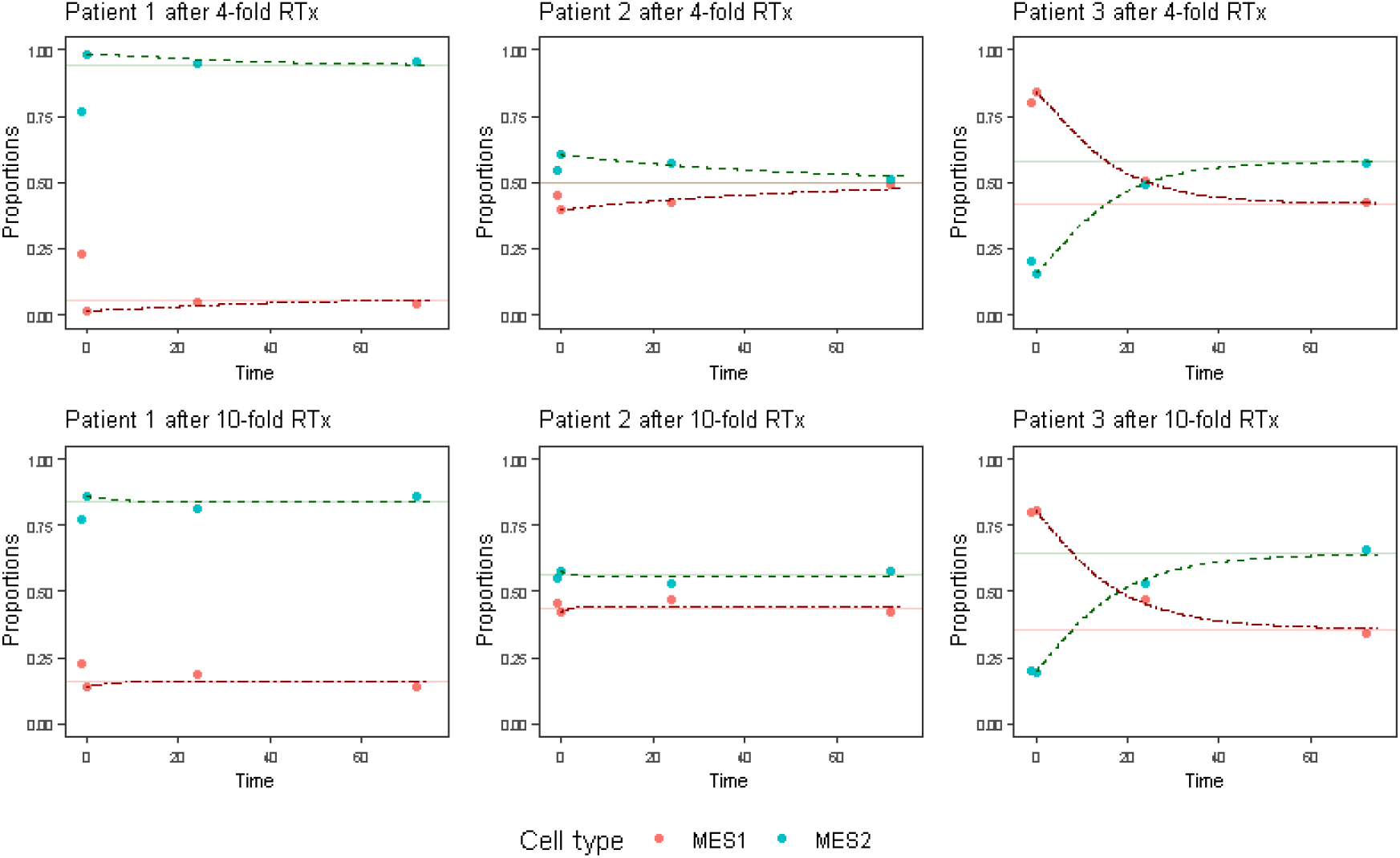
Fitted replicator dynamics equations vs. the measurements and ESS. First point (not fitted) indicates the measurement before the therapy is applied.

## 4 Discussion

Glioblastoma is a universally fatal disease for which there are no curative treatments and all patients eventually relapse after radiation and temozolomide chemotherapy. Treatment-induced resistance in cancer is thought to be caused by stochastic genetic or epigenetic driven phenotypes that preexist in small populations of tumour cells or evolve during the course of treatment [54, 55, 6, 13]. Tumor markers that predict the emergence of treatment refractory subpopulations in human tumors before and during the course of standard treatment may be exploited to adapt treatment and prevent relapse. Here we used machine-learning techniques to estimate how proportions of relevant glioblastoma cell types change over time under treatment from transcriptomics organoid data. We then fitted this data to the most common evolutionary game model, replicator dynamics, to predict long-term response to irradiation in these organoids.

To identify different cell subpopulations, we utilized both unsupervised and supervised machine learning methods. The unsupervised methods are attractive to researchers as they do not require any assumptions about the number of cell types present in the mixture, nor the profiles of said types. However, this makes them trickier to work with and less immediately interpretable. To overcome this shortcoming in interpretability, we selected genes with a high-fold change between their values in different profiles and compared them to both genes seen in signatures of published single cell analyses and genes from known pathways in standard pathway databases. Perhaps unsurprisingly, we found that the patient tumor samples were deconvoluted into immune cells, cycling cancer cells and healthy neuronal cells. This led to issues extrapolating from tumor profiles obtained *in vivo* to those *in vitro*, as not all cell types seen *in vivo* will be expected to be present in the *in vitro* situation. In the patient-derived organoids, for instance, immune cells, neurons, and blood vessels are missing.

Another issue with the unsupervised methods was the need to determine the number of cell types present *a priori*. If no assumptions about this are made, then extra steps are needed to estimate the optimal number of types empirically from the data. In our experience, both in this work and earlier unpublished work, this step does not generally yield a clear result when working with real datasets.

We found the supervised methods more interpretable, as they immediately allowed us to extract and interpret key information about the expected cell types present in the mixture; a key advantage over the unsupervised methods.

However, there are issues extrapolating from single cell data to bulk data, as the difference in measurement platforms can cause systematic differences in measurement that would be seen in the cell profiles (similarly to batch effects between datasets measured on different platforms).

Further, the supervised analysis relies on there being appropriate single-cell datasets available for training, something which will not be the case for all scenarios where deconvolution results would be useful.

Another limitation of this study is that tumors have spatial aspects, for instance hypoxic cells are more common in the center/core of the tumor where the supply of oxygen is limited (diffusion-limited hypoxia) or in the neighborhood of defective vessels (acute hypoxia). The spatial aspects could be analysed through single-cell imaging data, for instance [56], or through taking separate samples for bulk analysis from the core and the rim [57] and then using spatial game-theoretic models to predict further spatio-temporal response. Cells could then be assumed to only interact with the cells in their local neighbourhood, so local dynamics would become important, and tumors are not assumed to be well-mixed [20]. In case of using a replicator dynamics model, one could assume that different regions correspond to different fitness matrices, as done in [58], because cells’ chances for proliferation and survival will vary with conditions in their neighborhood.

Our results from the supervised analysis suggest that all three of our analysed organoids contain mainly mesenchymal cells, with some being the MES-like 2 hypoxia-dependent cells and others being the MES-like 1 hypoxia-independent cells. As the cell profiles we trained on were based on Neftel’s data [35] they can be directly compared to the results from Shakya et al [57] where they showed that the hypoxic organoid core mimicked the MES-like 2 profile. However, in their work they saw that the other cells in the organoid, in particular the cells in the organoid rim, expressed genes from the AC-like or OPC-like profiles. One key difference between their work and our approach is that we directly train a predictor of the cell type proportions from their single cell data, whereas they use the meta-modules of genes seen to be consistently differentially-expressed in these cells, this difference could potentially lead to these anomalies and should be investigated further.

Further it should be noted, that the decrease in the hypoxia-independent cells in response to radiation seen in our results for patient 3 is expected, as radiation-resistance is known to be related to hypoxia e.g. [59] and therefore the hypoxia-dependent cells (MES-like 2) can be expected to be more radio-resistant.

For prediction of the long-term response in organoids we fitted the identified proportions of different cell types to replicator dynamics [48, 49], which model classic and likely the most simple evolutionary dynamics. While the fit of data to replicator dynamics was very good, we could use different game-theoretic dynamics, such as Darwinian or adaptive dynamics [60, 61, 62]. As the total cell volume in organoids considered in the experiments carried out here was kept constant, to use these different dynamics we would need to assume that the population equilibrium was reached and fit only the evolutionary dynamics. Alternatively, we could combine volumetric data from the patients to fit the population dynamics, similarly as we did in [63, 64, 65], while fitting the evolutionary dynamics that are very difficult to estimate *in vivo* from the patient-derived organoids. An advantage of this approach could be that we would not need to assume that there is a fixed number of possible cell types with a fixed profile, but instead could trace radioresistance as an evolving trait, similarly to others modeling evolving traits responding to treatment [13, 66, 67]. Once the used model is validated, we could optimize the treatment, for example using the Stackelberg evolutionary game framework [66, 68, 69].

While our game-theoretic models predicted coexistence of the two different tumor types in organoids, we did not collect additional measurements to validate whether these predictions are correct. This was also not the main goal of our study; rather we wanted to demonstrates all steps needed for utilization of transcriptomics data in predictive game-theoretic modeling. Validation of our predictions and comparing the fit and predictive capabilities of different game-theoretic dynamics will be our natural next step. More time series data will be collected to perform both data fitting and validation. Once we have a validated game-theoretic model of the radiotherapy response in organoids, we can predict the impact of alternative treatments (including their timing and dosing), in order to delay/predict therapy resistance in the corresponding patients. A huge advantage of organoids is that these predictions can be then validated through real experiments. This is because in many studies organoids have been shown to retain major patient characteristics that determine treatment resistance [70, 71, 27, 28]. As such, we can utilize game-theoretic models validated through organoids to improve treatment in real patients.

Despite its limitations, this study shows that it is possible to apply both supervised and unsupervised deconvolution techniques to bulk gene expression data from tumors and to use the information obtained on the changes in cell type proportions over time to fit game-theoretic models of the tumor eco-evolutionary response to treatment. This is an important step forward in the development and application of these models and their use with real data predicting both ecological and evolutionary cancer dynamics and designing evolutionary therapies, i.e. therapies that anticipate and forestall treatment-induced resistance in cancer cells.

For our next steps we note that the TGCA dataset contains many different sets of measurements on the same patients. In this work we have only made use of the transcriptomics data, given all these measurements are made on the same tumors, which can be assumed to be composed of the same types, one direction for further research is learning to leverage the additional information available in other measurement sets from these tumors to improve the deconvolution overall and to improve the fit and predictive capabilities of game-theoretic models.

We believe that using cancer data in a mathematical model and adapting treatment strategies to the resulting predictions has the potential to revolutionise personalised medicine. When facing such a heterogeneous formation as cancer, one must apply a therapy adapted to the targeted malignancy. Our ultimate goal is to provide a process which, using the available information about the tumour composition and its measurements combined with a game-theoretic model, will simplify the selection of a better cancer treatment strategy.

## Acknowledgements

We would like to thank Dr. Jafar Rezaei for his valuable feedback that improved quality of this work. This research was supported by European Union’s Horizon 2020 research and innovation programme under the Marie Skłodowska-Curie grant agreement No 955708, the Dutch National Foundation projects ENWPR.020.006, and OCENW.KLEIN.277 and Dutch Cancer Society (KWF 11698) and funding from StopHersenTumoren.

